# Distribution of Panama’s narrow-range trees: are there hot-spots?

**DOI:** 10.1101/2021.02.13.431023

**Authors:** Elizabeth Tokarz, Richard Condit

## Abstract

**Background:** Tree species with narrow ranges are a conservation concern because heightened extinction risk accompanies their small populations. Assessing risks for these species is challenging, however, especially in tropical flora where their sparse populations seldom appear in traditional plots and inventories. Here, we utilize instead large scale databases that combine tree records from many sources to test hypotheses about where the narrow-range tree species of Panama are concentrated.

**Methods:** All individual records were collected from public databases, and the range size of each tree species found in Panama was estimated as a polygon around all its locations. Rare species were defined as those with ranges < 20,000 km^2^. We divided Panama into geographic regions and elevation zones and counted the number of individual records and the species richness in each, separating rare species from all other species.

**Results:** The proportion of rare species peaked at elevations above 2000 m, reaching 17.3% of the species recorded. At lower elevation across the country, the proportion was 6-11%, except in the dry Pacific region, where it was 1.5%. Wet forests of the Caribbean coast had 8.4% rare species, slightly higher than other regions. The total number of rare species, however, peaked at mid-elevation, not high elevation, because total species richness was highest there.

**Conclusions:** High elevation forests of west Panama have higher endemicity of trees than all low-elevation regions. Dry forests had the lowest endemicity. This supports the notion that montane forests of Central America should be a conservation focus, however, given generally higher diversity at low- to mid-elevation, lowlands are also important habitats for rare species.

## Introduction

Species with narrow ranges are vulnerable to extinction compared to widespread species due to small population size and narrow environmental tolerance (Hubbell *et al.*, 2008; Sykes *et al.*, 2020). Yet these narrow endemics can contribute substantially to local and regional diversity (Heegaard *et al.,* 2013). Maintaining their populations in the face of anthropogenic pressure requires knowing where they occur (Jenkins *et al.*, 2013). If many scarce species are concentrated in a particular region or habitat, then localized conservation efforts can cover multiple species (Brown *et al.*, 1995). But ‘hot-spots’ of rare species may not coincide with areas of high overall diversity (Prendergast *et al.*, 1993), so it is important to understand the distribution of narrow-range species separately from widespread species. In diverse tropical forests, ranges are seldom known completely, and aggregating across many rare species offers a way not only to identify hot-spots but also to improve inference in poorly-explored regions.

Panama is highly diverse due to its strong precipitation and soil gradients (Gentry, 1982, 1992; Condit *et al.*, 2004, 2011, 2013; Barthlott *et al.*, 2005), and it has many rare tree species (Hubbell & Foster, 1986; Condit *et al.*, 2020). Hot spots have been proposed, including the Choco region along the Caribbean of eastern Panama into Colombia, and the high mountains of western Panama into Costa Rica (Stattersfield *et al.*, 1998; Montanez & Angehr, 2007; Pérez-Escobar *et al.*, 2019; Fagua & Ramsey, 2019). Panama also has an isolated fragment of true dry forest with an unusual flora. Our goal is a rigorous assessment of the number of rare tree species relative to total diversity, and whether concentrations of rare species follow the proposed hot-spots.

We have established many tree plots in Panama (Condit, 1998; Condit *et al.*, 2002, 2013), but the rarest species seldom appear in plots (Condit *et al.*, 2020). Our assessment thus requires wider data collecting, only possible by databases that combine records from many observers (Maitner *et al.*, 2017). Using two large online sources, we completed full range estimates of all tree species known in Panama, verifying both the species and their coordinates using expert knowledge (Condit *et al.*, 2020). Here we define rare species as those with narrow ranges (Rabinowitz, 1981; Magurran, 2004; Espeland & Emam, 2011; Flather & Sieg, 2013), and we test the hypothesis that rarity thus defined is concentrated in certain regions, including the proposed hot-spots. We calculate endemicity using two definitions: 1) the proportion of rare species relative to all species, and 2) the total number of rare species.

## Methods

### Narrow-range species

A checklist of all tree species in Panama ≥ 3 m tall was thoroughly vetted, eliminating invalid names, harmonizing taxonomic synonyms, and confirming records within the country (Condit *et al.*, 2020). The full list is available in Condit *et al.* (2019). All records with coordinates for those confirmed species were then collected from public, online sources at BIEN (Maitner *et al.*, 2017) and the Missouri Botanical Garden (http://legacy.tropicos.org/Home.aspx). Duplicate sets of coordinates within any species were discarded. Further details can be found in Condit *et al.* (2020).

The global range size – Panama and elsewhere – of a species was estimated as the area of the minimum convex polygon enclosing all occurrences, excluding large bodies of water (Condit *et al.*, 2020). There were 448 species with at least three sets of coordinates that had a range < 20,000 km^2^, plus 46 more species having 1-2 records (so a tiny range that could not be calculated). We define those 494 species as rare and the remaining 2549 species as common, so across Panama, 16.3% of all species were in the rare category. Many of the species occur outside Panama, even the rare ones, but our question here is where they occur within Panama.

### Elevation

We downloaded shuttle radar topography at 90-m resolution from the CGIAR Spatial Data Consortium (http://srtm.csi.cgiar.org/srtmdata/). These were interpolated using the R raster package (https://rspatial.org/raster/) to estimate the elevation of every tree record in Panama. Our first question is whether rare species are concentrated at certain elevations, so we divided Panama into bands of 250 m vertical range from sea level to 1500 m ASL, one band 1500-2000 m, and a final band > 2000 m, and we counted the number of rare and common species in each band. We also estimated the sample size per elevation band as the number of distinct records (eliminating duplicate coordinates within species) for rare and common species (Table 1).

**Table 1:**
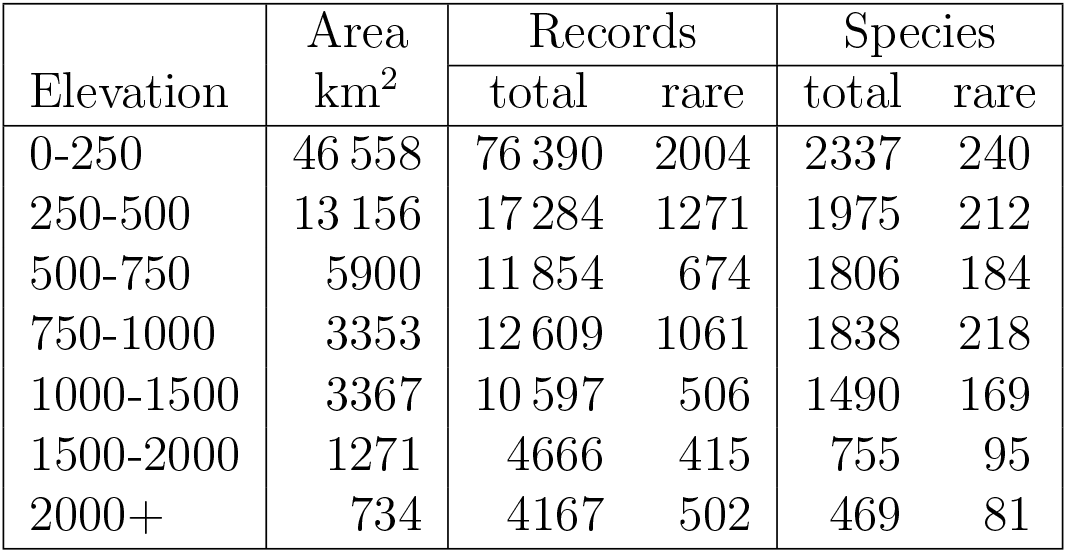
Area, collection effort, and observed species richness of Panama across elevation zones. Rare species are defined as those with ranges < 20, 000 km^2^.

### Climatic provinces

Besides elevation, the important habitat division in Panama is due to rainfall, and the country is divided into a relatively dry Pacific slope and a wet Caribbean slope (Condit *et al.*, 2002, 2004). The mountain chain running the length of the isthmus creates a division between the two slopes (Condit *et al.*, 2011). The Caribbean slope is wet forest according to the Holdridge system (Holdridge, 1967), while the Pacific slope is variable, including dry and moist forest with some pockets of wet. True Holdridge dry forest occurs only in the ‘arco seco’ on the east side of the Azuero Peninsula (Fig. 1).

**Figure 1:**
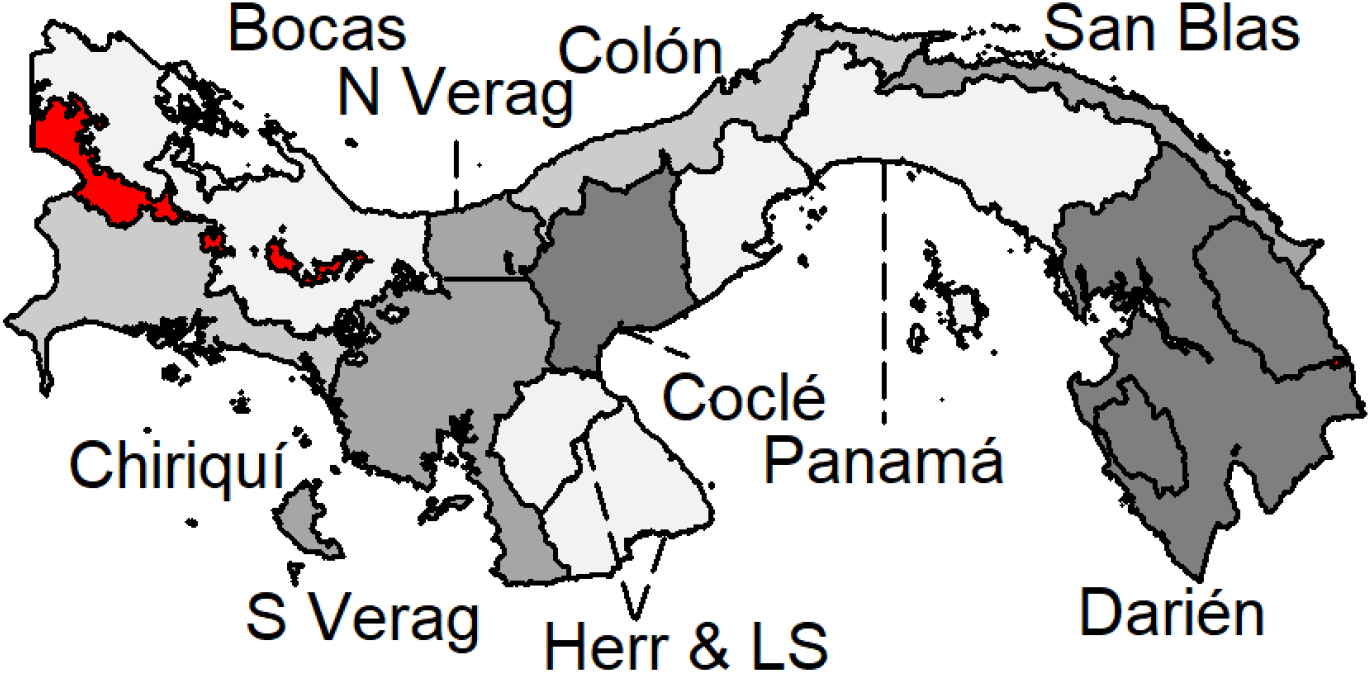
Panama province map. The high elevation ‘province’ (> 1500 m) is red. Abbreviated names: Bocas=Bocas del Toro; N Verag=North Veraguas (north of 8.51° N.), S Verag=South Veraguas (the Veraguas sections are colored the same), Her-LS=Herrera and Los Santos (combined in analyes; the line within demarcates the two provinces). Two enclosed areas within Darien are indigenous comarcas, considered part of Darien in analyses. The line within Panama province is the Canal, not a political division.

To evaluate whether rare species occur disproportionately relative to this division between Pacific and Caribbean slopes, we took advantage of Panama’s political provinces, most of which are either on the Caribbean or the Pacific slope. The province of Veraguas is one exception, so we split it into two at 8.51° N. latitude, creating North Veraguas on the Caribbean and South Veraguas on the Pacific. The province of Darien also crosses the isthmus, but it was not divided because the mountain spine does not continue that far east, and the wet-dry contrast breaks down. Finally, the provinces Herrera and Los Santos were combined because they are small and adjacent, and they are exclusively dry forest (Fig. 1). We utilize the old province borders of Panama, prior to demarcation of Native American comarcas, because the original borders split the Caribbean and Pacific slopes better.

In most of Panama, the central mountain ridge dividing Caribbean and Pacific is < 1000 m elevation. In the far west, however, elevations are higher, including a region > 1500 m, up to 3400 m, in two western provinces (Fig. 1). We thus created an additional high-elevation ‘province’ spanning Bocas del Toro and Chiriqui, leaving the lowlands of Chiriqui on the Pacific slope and Bocas del Toro on the Caribbean slope. The high-elevation province combines the two highest elevation bands (Table 1). As for elevation bands, the number of species and number of records in rare and common groups were counted in each province (Table 2).

**Table 2:** Area, collection effort, and observed species richness of Panama across provinces. Herr & LS is the combined Herrera and Los Santos provinces. High elev is the zone ≥ 1500 m elevation, which combines the two highest elevation bands (Table 1). Rare species are defined as those with ranges < 20, 000 km^2^.

| Province | Area<br>km<sup>2</sup> | Records |  | Species |  |
| --- | --- | --- | --- | --- | --- |
|  |  | total | rare | total | rare |
| Bocas | 11 196 | 6657 | 216 | 1206 | 78 |
| Chiriqui | 6335 | 10 938 | 426 | 1466 | 124 |
| Cocle | 4831 | 9357 | 710 | 1152 | 85 |
| Colon | 4466 | 26 862 | 1083 | 1552 | 131 |
| Darien | 15 833 | 13 087 | 325 | 1427 | 71 |
| Herr & LS | 6023 | 2280 | 21 | 569 | 9 |
| Panama | 11 018 | 46 067 | 1992 | 1871 | 180 |
| San Blas | 2302 | 7542 | 570 | 1169 | 103 |
| N Veraguas | 1744 | 3205 | 193 | 848 | 51 |
| S Veraguas | 8589 | 4300 | 108 | 967 | 47 |
| High elev | 2005 | 8835 | 917 | 850 | 119 |

### Rarefaction

The sampling effort and the surface area in the different elevation bands, and in different provinces, varied considerably, so a direct comparison of species counts in the units is biased. By rarefying to a fixed sample size in each elevation band, or province, we accounted for differences in land area as well as variation in collection effort.

The elevation zone > 2000 m had the fewest occurrences (4167, all species combined), so that was set as the rarefaction sample for elevation comparisons. In each zone, we randomly extracted 4167 occurrences, without replacement, 1000 different times. For rarefaction of province occurrences, we chose a sample of 3205, which was the second smallest; we avoided the smallest, 2280 occurrences in Herrera-Los Santos, because it was so much lower. We also repeated the province rarefaction with the smaller sample, and the results were unaffected, so we do not report it.

Species-accumulation with sample size was also tested to examine the magnitude of biases in collection effort. Across the entire nation of Panama, random draws were made using sample sizes

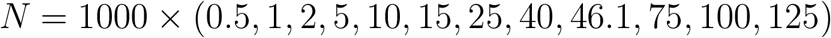

occurrences, with 1000 replicates at each sample size. This creates an accumulation curve, showing how the number of common and rare species, and their ratio, increased with sample size. We also created an accumulation curve from within one of the provinces, using 1000 replicates of the same samples up to 46.1 × 10^3^ km^2^, the latter the entire province (Table 1). We chose the province with the most samples, the one named Panama (not to be confused with the identical country name).

### Identifying hot spots for rare species

In order to identify hot spots for rare tree species in Panama, we used two test statistics. One is simply the number of rare species in every sampling unit, based on rarefaction to a fixed sample. The second is the ratio of rare species to all species, again from a rarefied sample. The 1000 replicates of rarefied samples produce a standard deviation, which we report as the standard error (SE) of each estimate of rarefied species richness and rarefied proportion rare species. In those cases where the rarefied sample size ≤ the full sample (two provinces, one elevation zone), rarefaction was carried out with replacement to estimate SE; otherwise, sampling was without replacement. We interpret ±2×SE as 95% confidence limits, and consider estimates with non-overlapping confidence intervals to be statistically distinct.

### Data available

A supplementary data archive includes the full species list with range sizes available for download (Condit *et al.*, 2019).

## Results

### Elevation

The ratio of rare species to all species increased with elevation, from 5.5% in the lowest elevation band to 17.3% at the highest (based on fixed sample sizes for comparison) (Table 3). Based on confidence limits of (2×SE), the percentages at high elevation and at low elevation were statistically distinct from the remaining elevation zones (Table 3).

**Table 3:** Number of total tree species, number of rare species (those with ranges < 20, 000 km^2^), and the percent rare in elevation zones of Panama, calculated from subsamples of 4167 records in each zone, with standard errors (SE) among replicates of the rarefied subsamples.

| Elevation | All species |  | Rare species |  | Percent rare |  |
| --- | --- | --- | --- | --- | --- | --- |
|  | total | SE | total | SE | mean | SE |
| 0-250 | 1181.7 | 15.9 | 65.2 | 5.7 | 5.5 | 0.46 |
| 250-500 | 1346.8 | 14.8 | 133.8 | 6.1 | 9.9 | 0.44 |
| 500-750 | 1327.4 | 14.9 | 119.1 | 5.8 | 9.0 | 0.41 |
| 750-1000 | 1332.1 | 14.3 | 143.5 | 5.8 | 10.8 | 0.41 |
| 1000-1500 | 1154.2 | 12.8 | 117.6 | 5.4 | 10.2 | 0.43 |
| 1500-2000 | 733.1 | 4.5 | 92.9 | 1.4 | 12.7 | 0.19 |
| 2000+ | 469.0 | 0.0 | 81.0 | 0.0 | 17.3 | 0.00 |

The total number of species (common plus rare) in fixed samples was similar from 0-1500 m, then dropped abruptly and significantly at elevations above 1500 m. There were thus two opposing trends with increasing elevation - fewer total species but a higher percentage rare, and this meant that the total number of rare species peaked at mid-elevation, in the 750-1000 m zone (Table 3).

### Province

The ratio of rare species to all species was very low in Herrera/Los Santos, the driest zone (1.5%), and it was highest in the montane ‘province’ (14.9%). Both those extremes were statistically distinct from the remaining provinces. Other provinces had 4-9% rare species (Table 4). The total number of rare species in fixed samples was high in San Blas and the high-elevation province, and also in lowland Chiriqui (< 1500 m). The number of rare species was significantly lower in several provinces, especially so in Herrera/Los Santos. The total number of species (common and rare) in fixed samples was 850-1050 in most provinces, but significantly lower in the high-elevation province and much lower in Herrera/Los Santos (Table 4).

**Table 4:** Number of total tree species, number of rare species (those with ranges < 20, 000 km^2^), and the percent rare in Panama provinces, calculated from subsamples of 3205 records in each province, with standard errors (SE) among replicates of the rarefied subsamples. In Herrera-Los Santos (Herr & LS), the subsample was only 2280.

| Province | All species |  | Rare species |  | Percent rare |  |
| --- | --- | --- | --- | --- | --- | --- |
|  | total | SE | total | SE | mean | SE |
| Bocas | 989.5 | 10.9 | 56.5 | 3.4 | 5.7 | 0.33 |
| Chiriqui | 1038.7 | 12.8 | 71.1 | 4.8 | 6.8 | 0.44 |
| Cocle | 902.6 | 10.9 | 65.9 | 3.1 | 7.3 | 0.34 |
| Colon | 985.1 | 13.3 | 63.0 | 4.6 | 6.4 | 0.45 |
| Darien | 1028.4 | 12.6 | 42.4 | 3.7 | 4.1 | 0.34 |
| Herr & LS | 520.1 | 6.0 | 7.7 | 1.0 | 1.5 | 0.19 |
| Panama | 1031.6 | 14.7 | 76.3 | 5.7 | 7.4 | 0.52 |
| San Blas | 959.3 | 10.2 | 80.9 | 3.4 | 8.4 | 0.35 |
| N Veraguas | 848.0 | 8.8 | 51.0 | 2.4 | 6.0 | 0.33 |
| S Veraguas | 883.0 | 7.8 | 39.6 | 2.4 | 4.5 | 0.26 |
| High elev | 637.9 | 10.2 | 94.8 | 3.9 | 14.9 | 0.61 |

### Rarefaction

The total number of species, or rare species, increased steeply with sample size with < 10,000 records, but reached a plateau beyond 25,000 (Fig. 2).

The ratio of rare to total species also increased steeply in small samples, but rose much more slowly beyond 25,000, and finally reaching 16.3% in the entire country. Within a single province, the proportion of rare species rose quickly in small samples, but stabilized at 9.5%, well below the proportion in a sample of the same size drawn from the entire country (Fig. 2).

**Figure 2:**
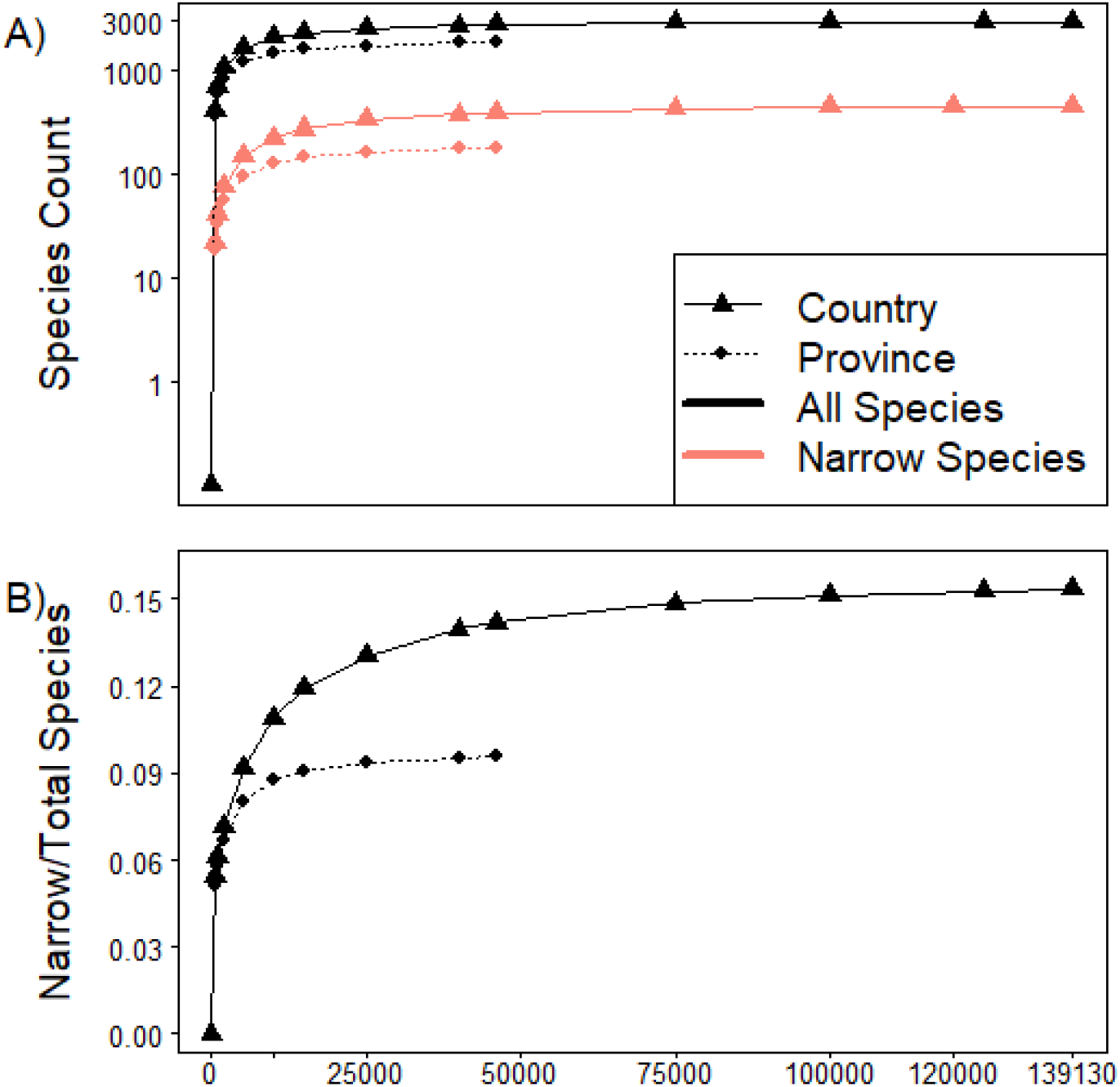
Rarefaction. Each panel shows species counts or ratios in randomly drawn subsamples of various sizes drawn from all records (see Methods). A) Total number of species (black) and rare species (red). Solid lines are results when samples were taken from the entire nation of Panama; dashed lines from within one province (the province named Panama). B) The ratio of rare species to total species. The solid line is from the entire country, the dashed from within the single province named Panama.

## Discussion

Defining hot-spots using the proportion of narrow-range tree species relative to all species, we found that the highest mountains of Panama, > 1500 m elevation, stood out. The proportion of rare species was much higher there than the rest of the country. The Caribbean slope of eastern Panama, in the Choco biogeographic province, did not stand out by this measure. The dry forests of Panama had the lowest proportion of narrow endemics.

The results changed, however, when counting the total richness of narrow-range tree species, rather than their proportion. The total number of rare species at high elevation was similar to the total number in San Blas Province of the Choco region, and other provinces were only slightly behind. This was possible simply because the montane region had far fewer species in total. Dry forests of Herrera-Los Santos provinces, on the other hand, had few rare species whether counting the total number or their proportion.

Previous studies have reported higher proportions of rare or endemic species at high elevations in the Neotropics (Zizka *et al.*, 2018; Knapp, 2002) and globally (Steinbauer *et al.*, 2016). At the same time, high elevation forests in the Neotropics have fewer plant species (Gentry, 1988), and there is a global pattern of diversity peaking at mid-elevations (McCain, 2005; Rahbek, 1995; Beck *et al.*, 2017). Our results parallel both these findings. It is not clear, however, that anyone has put the two patterns together: a higher proportion of endemics but fewer total species at high elevation means, in our analysis, that there were only slightly more narrow endemics in the high elevation zone of Panama than in other areas.

High endemism and overall richness of the Choco region, including the Caribbean forests of San Blas Province in Panama, is widely quoted (Fagua & Ramsey, 2019; Pérez-Escobar *et al.*, 2019). Our work in tree plots shows high local diversity in these wet forests of the Caribbean (Condit *et al.*, 2005; Pérez *et al.*, 2005), but this does not address total richness (gamma diversity) or endemism. Our results here do not favor unusually high endemism in San Blas, as the proportion of rare species there was only slightly higher than the rest of Panama.

The dry forest zone of Panama is small, and it harbors unusual species for a wet tropical climate. Example of trees restricted to the dry zone include *Prosopis juliflora* (the mesquite, Fabaceae) and *Gyrocarpus americana* (Hernandiaceae). Their populations within Panama are small and isolated. Though locally rare, however, these species have very wide ranges across the Neotropics; *Prosopis* occurs from the southern USA to Chile. We found tree richness and endemism far lower in the dry forest zone of Panama than anywhere else in the country. Our plot inventories also show lower tree diversity in dry areas (Pérez *et al.*, 2005).

## Conclusion

Narrowly distributed tree species were formed a higher proportion of the tree diversity at elevations greater than > 1500 m. At lower elevation and across the rest of Panama, the proportion of these narrow endemics was lower. This favors conservation efforts directed toward high elevation habitats. Our study joins many others that highlight the importance of high elevation habitats for biodiversity (Körner, 2004; Nogué *et al.*, 2012; Rahbek *et al.*, 2019).

At the same time, our results show higher overall species richness at low and mid-elevations. Hence, the gradient in total richness and the gradient in percentage endemics are opposite, so the total richness of rare species (not the proportion) is similar across the elevation gradient. The fact that low or mid-elevation forests have the highest species richness is well known, and conservation efforts in low elevation forests, despite a low percentage endemism, is certainly as important as efforts directed toward the small areas at high elevation.

## Declarations

### Ethics approval and consent to participate

The research involved no human subjects. Permission from owners was always obtained for working in parks or on private land in Panama.

### Consent for publication

Not applicable.

### Availability of data and material

A supplementary data archive shows additional results (Condit *et al.*, 2019). All species names, synonyms, and the full data, including all individual records and estimated range sizes, are available there for download.

### Authors’ contributions

ET wrote the manuscript. RC directed plot censuses and inventories, assembled floristic records, and calculated range sizes.

## Acknowledgements

We thank the staff of the Center for Tree Science at the Morton Arboretum for financial and scientific support.

